# Large-scale RNA editing profiling in different adult chicken tissues

**DOI:** 10.1101/319871

**Authors:** Hamid Shafiei, Mohammad Reza Bakhtiarizadeh, Abdolreza Salehi

## Abstract

RNA editing is a post-transcription maturation process that diversifies genomically encoded information and can lead to diversity and complexity of transcriptome, especially in the brain. Thanks to next-generation sequencing technologies, a large number of editing sites have been identified in different species, especially in human, mouse and rat. While this mechanism is well described in mammals, only a few studies have been performed in the chicken. Here, we developed a rigorous computational strategy to identify RNA editing sites in eight different tissues of the chicken (brain, spleen, colon, lung, kidney, heart, testes and liver), based on RNA sequencing data alone. We identified 68 A-to-G editing sites in 46 genes. Only two of these were previously reported in chicken. We found no C-to-U sites, attesting the lack of this type of editing mechanism in the chicken. Similar to mammals, the editing sites were enriched in non-coding regions, rarely resulted in change of amino acids, showed a critical role in nervous system and had a low guanosine level upstream of the editing site and some enrichment downstream from the site. Moreover, in contrast to mammals, editing sites were weakly enriched in interspersed repeats and the frequency and editing ratio of non-synonymous sites were higher than those of synonymous sites.

Interestingly, we found several tissue-specific edited genes including GABRA3, SORL1 and HTR1D in brain and RYR2 and FHOD3 in heart that were associated with functional processes relevant to the corresponding tissue. This finding highlighted the importance of the RNA editing in several chicken tissues, especially the brain. This study extends our understanding of RNA editing in chicken tissues and establish a foundation for further exploration of this process.

## Introduction

There are many biological processes that contribute to transcriptome diversity, such as alternative splicing, polyadenylation and RNA editing. These processes can result in the diversification of transcriptome as well as protein repertoire (Nishikura, 2010). RNA editing is a process of RNA maturation and refers to the modification of specific RNA nucleotides in the co-transcriptional and post-transcriptional stages of gene expression, without altering its template genomic DNA. This mechanism can flexibly and dynamically increase the diversity of expressed transcripts as well as proteins, beyond the genomic blueprint (Li and Church, 2013). The most prevalent RNA editing substitution in mammals is enzymatic deamination of adenosine nucleotides to inosine (recognize as guanosine (G) during translation by the cellular machinery and reverse transcription), by adenosine deaminase acting on RNA (ADAR) family enzymes (Huntley *et al*., 2016). There are three members of ADAR family in vertebrates, comprising ADAR1 (ADAR), ADAR2 (ADARB1) and ADAR3 (ADARB2) (Gu *et al*., 2012). While ADAR1 and ADAR2 are expressed in almost all human tissues and have overlapping target specificities, ADAR3 is only expressed in the brain without a known A-to-I RNA editing activity. While there is no proof to support the enzymatic activity of ADAR3, it contains conserved functional domain specification (Savva *et al*., 2012). These enzymes are present in nucleus and cytoplasm, however, ADAR2 is mainly a nuclear enzyme. Another chemical change of specific RNA residue is cytosine to uridine (recognize as thymine (T) during translation by the cellular machinery and reverse transcription) substitution that is catalyzed by the member of activation-induced deaminase/apolipoprotein B editing complex catalytic (AID/APOBEC) family (Bahn *et al*., 2012, Liu *et al*., 2014). This type of editing event is a very rare mechanism in the mammalian transcriptome and to date only a few sites have been discovered (Gu *et al*., 2012, Huntley *et al*., 2016).

Previous studies have reported potential roles of RNA editing in different aspects of gene expression including splicing (Nishikura, 2010), microRNA targeting (Nishikura, 2016), RNA localization (Chen and Carmichael, 2009), RNA stability and degradation (Stellos *et al*., 2016), translation and other important cellular processes (Nishikura, 2016). Also due to the high frequency of RNA editing in the brain transcriptome, it has been known as a key factor in the development of higher brain functions (Peng *et al*., 2012b, Li and Church, 2013). Moreover, dysregulation of RNA editing might be involved in various diseases; such as tumor malignancy, dyschromatosis symmetrical hereditary, sporadic amyotrophic lateral sclerosis (ALS), neuropsychiatric disorders, autism, and cancer that emphasize the biological significance of this mechanism (Li and Church, 2013, Huntley *et al*., 2016).

The development of high-throughput sequencing technology such as RNA-Seq method and the open access data sharing in genomic research have facilitated the discovery of RNA editing events and its functional mechanisms in different organisms. Hence, many genome-wide characterizations of RNA editing studies have been performed in primate such as human (Bahn *et al*., 2012, Li and Church, 2013, Bazak *et al*., 2014, Mo *et al*., 2014, Picardi *et al*., 2015), mouse (Danecek *et al*., 2012, Gu *et al*., 2012, Lagarrigue *et al*., 2013, Huntley *et al*., 2016), cattle (Pinto *et al*., 2014, Bakhtiarizadeh *et al*., 2018), pig (Larsen *et al*., 2016), chicken (Frésard *et al*., 2015, Roux *et al*., 2016) using this method, that has expedited the exploration of a large number of RNA editing sites (Picardi *et al*., 2015). The importance of chickens in evolutionary biology is quite obvious, as they are known as an ideal organism for filling the evolution gap between mammals and other vertebrates (Roux *et al*., 2016). In our best knowledge, only three studies have been done for genome-wide identification of RNA editing in chicken. In one of these studies, 36 A-to-G RNA editing sites were reported in brain, heart and liver tissues of chicken embryos (Frésard *et al*., 2015). In another study, 25 RNA editing sites were identified in liver and abdominal adipose tissue, of which three events had been previously characterized in mammals (Roux *et al*., 2016). Also, Hung et. al., applied RNA-Seq data from 20 metazoan species (including chicken), to identify high confidence editing sites. In chicken, 94 A-to-G editing sites were reported, which distribution of these sites had not been discussed in different tissues (Hung *et al*., 2017). Therefore, in spite of the biological importance of RNA editing, a few studies have been performed in chicken, which indicate much more work is needed to identify the extent of this mechanism in this species. On the other hand, there has been no comprehensive study investigating editome in many different chicken tissues. RNA editome profiling in chicken tissues may shed light on novel physiological roles of RNA editing in this species. Hence, to achieve this goal and characterize the complexity of RNA editing in different chicken tissues, we performed a genome-wide study using strand-specific RNA-Seq data across eight tissues from three adult chickens. We developed a computational strategy to find potential editing events in chicken using RNA-Seq data alone.

## Material and Methods

### RNA-Seq datasets and preprocessing

In order to investigate the RNA editing sites in chicken, we downloaded 22 publicly available paired-end RNA-Seq samples related to eight tissue types including brain, colon, heart, kidney, liver, lung, spleen and testes from three adult chickens (NCBI GEO accession GSE41637). The original data and samples details are described in (Merkin et al., 2012). This dataset includes strand-specific poly(A)+ RNA sequencing from different tissues of five species (three individual per species). cDNA libraries of these samples were sequenced on an Illumina HiSeq 2000 platform. Here, we focused only on chicken samples. In this dataset, samples of a chicken were sequenced to high coverage (876.2 million reads per sample, on average, with length 76 bp), while the two other chicken samples had moderate coverage (382.4 million reads per sample, on average, with length 36 bp). The qualities of the raw reads were checked using FastQC v0.11.5 (Schmieder and Edwards, 2011). Then, raw reads were subjected to adapter removal and quality trimming prior to alignment to the reference genome using Trimmomatic program (Bolger *et al*., 2014) (parameters of Trailing 20 Maxinfo 60:0.90, and Minimum length 60 for 76 bp and Trailing 20 Maxinfo 30:0.90, and Minimum length 30 for 36 bp datasets).

### Mapping and variant calling

The cleaned reads were aligned to reference genome (Gallus gallus 5.0) using HISAT2 aligner (Liang *et al*., 2016). To improve the alignment of RNA-Seq reads, we inputted exon-intron junction information into the aligner using the annotation GTF file (Ensembl version 88). Subsequently, uniquely and concordantly mapped reads for each sample were extracted by Picard tool for further analysis. Furthermore, PCR-induced duplications were marked and excluded from analysis using MarkDuplicates tool from the Picard package (Choi *et al*., 2015). Next, we proceeded to the variant calling step by realigning (to promote the aligning in the flanking of insertion and deletion regions) and recalibrating (to improve the quality of reads) the alignments with Genome Analysis Toolkit (GATK version 3.5) (Puckelwartz *et al*., 2014). In the next step, single nucleotide variants (SNVs) were first called using HaplotypeCaller method from GATK tool with a stand_call_conf and stand_emit_conf value of 30 and mbq of 25. To increase the confidence of identifying true SNVs, the variants were filtered by the GATK standard filters (Total depth of coverage <10, HomopolymerRun >5, RMSMappingQuality <40, MappingQualityRankSum < −12.5 QualitybyDepth < 2 and ReadPosRankSum <-8). To remove possibly false positive RNA editing sites due to SNPs, the remaining SNVs were filtered against known chicken SNPs found in Ensembl chicken SNP database version 146. We also discarded SNVs by several quality-aware filters including sites with more than one non-reference types, homozygous sites for the alternative allele, sites with <3 reads supporting the SNV and sites with extreme or rare degree of variation (threshold for the editing ratio was between 10% and 90%). To filter spurious SNVs arising from regions of the genome that are problematic for short read alignment, we also filtered SNVs which were located in regions with bidirectional transcription, simple sequence repeat (SSR) and 5 bp from a splicing site. To do this, bidirectional and splice site positions were obtained from annotation GTF file of chicken. Also, regions containing SSRs in chicken genome were identified by using GMATo software (Wang *et al*., 2013) (SSRs ranging in length from 1 to 8 nucleotides with a minimum length of 6 bases). For added stringency and to reduce false positives SNVs that arose through misalignment of sequencing reads to regions highly similar to other parts of the genome, we performed a realignment filtering using BLAT tool (Kent, 2002). Hence, 100bp of flanking sequence (50bp upstream and 50bp downstream of the SNV) were retrieved and mapped against reference genome. Only the SNVs which were located in uniquely mapped sequences were retained for further analysis.

### RNA-editing detection

After all the filtering above, we obtained a collection of high confidence SNVs, which can be combination of SNPs and RNA editing sites. In order to distinguish the RNA editing sites from SNPs, we used a rigorous method (GIREMI tool), which uses allelic linkage between SNVs and calculate the mutual information of the mismatch pairs identified in the RNA-Seq reads to distinguish RNA editing sites and SNPs without requiring sample-specific genome sequence data. Then, GIREMI improves the predictive power with generalized linear models (Zhang and Xiao, 2015). To do this, GIREMI requires only alignment file, reference genome and a list of known SNPs for interested species (here, Ensembl chicken SNP database version 146 was used). It is well known that true editing sites are often present in different individuals, whereas low frequency SNPs are most likely not (Ramaswami *et al*., 2013). Therefore, the SNVs detected as RNA editing candidates by GIREMI were further filtered out and these candidates were considered high quality only if identified in at least two chickens. A bash Script enabled us to extract the strand orientation of the variant to identify A to G modifications. The editing ratio for each editing site was estimated as the percentage of reads with the A-to-G variant compared with all the reads covering that site.

### Sequence preferences analysis

WebLogo program was applied to build a consensus sequence logo and investigate the sequence context flanking of the identified A-to-G editing sites (Crooks *et al*., 2004). To do this, we analyzed the upstream and downstream 10-bases sequence flanking editing sites.

### RNA-editing sites annotation

We used SnpEff program (v 4.3), a variant annotation and effect prediction tool (Cingolani et al., 2012), to functionally annotate all candidate RNA editing sites and their potential mutational effects on associated genes. The gene set used in annotation was Ensembl version 88. We then used Enrichr (Chen *et al*., 2013) to reveal more information about the biological functions and pathways that are significantly enriched in edited genes by focusing on gene ontology (GO) terms (BP, biological process) and KEGG pathways (using a standard false discovery rate (FDR)<=0.05).

We also screened the chicken genome to investigate whether candidate editing sites were enriched in the interspersed repeats, such as long interspersed nucleotide elements (LINE) and short interspersed nucleotide elements (SINE) families, or not. To this end, the positions of all interspersed repeats sequences in chicken genome were determined using RepeatMasker tool (UCSC database [http://genome.ucsc.edu] (Tarailo-Graovac and Chen, 2009)).

### EST analysis

Expressed sequence tags (ESTs) had been considered as a valuable source in gene discovery, genomic sequence annotation and other functional genomic projects like gene expression profiling, before the appearance of deep sequencing technology such as RNA-seq (Bakhtiarizadeh *et al*., 2011, Bakhtiarizadeh *et al*., 2012). Here, we used publicly available chicken ESTs (from NCBI database) to further validate the identified A-to-G editing sites. To do this, flanking sequences of RNA editing sites with a length of 50 nt were used as queries for BLAST search (e-values < 10-5) against chicken ESTs. Then, ESTs with editing sites were counted for each candidate sites by an in-house script.

### Gene expression quantification

To quantify the transcript abundance, Salmon software (version 0.8.1), with sequence and GC bias correction enabled and k=25 for indexing, was used (Patro et al., 2017). Read counts for each transcript were normalized for transcript length and sequencing depth, using the transcripts per kilobase million (TPM) method.

Supplemental files available at FigShare. File S1 contains summary statistics for reads and alignment information of different samples. File S2 contains final list of all identified DNA-RNA differences in different tissues of bovine. File S3 contains list of tissue specific edited genes in different tissues of chicken. Also, editing ratio of each editing site in each tissue is provided. The marked sites are tissue-specific edited sites in related tissue. File S4 contains features of all the identified A-to-G editing sites in chicken transcriptome. File S5 contains Spearman correlation between gene expression of edited genes and their editing levels in different tissues. File S6 contains gene expression of the tissue specific edited genes and their editing levels in different tissues. File S7 contains comparisons of our results with the other studies in chickens.

## Results

### RNA editing detection

To construct a chicken reference atlas of editing events, we examined the RNA editome across eight tissues using 22 strand-specific RNA-Seq samples derived from three chickens (brain and liver tissues had two biological repeats). To this end, we developed a computational pipeline by using a rigorous strategy, which is summarized in Figure 1. Our method allowed to identify the editing sites directly from RNA-Seq data, without need for available matched DNA information from the same sample. The performance and efficiency of such method have been reported in previous studies (Ramaswami *et al*., 2013, Zhu *et al*., 2013, Han *et al*., 2015, Moran *et al*., 2017). In brief, RNA sequencing reads were first evaluated with FASTQC and processed to remove adaptors and low-quality reads using Trimmomatic. Then, clean reads were aligned to the reference genome Gallgall5 using Hisat2 and uniquely and concordantly aligned reads were extracted by Picard tool. Next, GATK tool along with a series of filters (to remove the noises), was used to call SNVs. After using BLAT software, to mitigate the effect of misalignments, we used GIREMI software, which is developed to detect RNA-editing sites from RNA-Seq data alone, to identify candidate editing sites. Finally, we called a SNV as a strong candidate if it appeared in at least two biological replicates.

**Figure 1.**
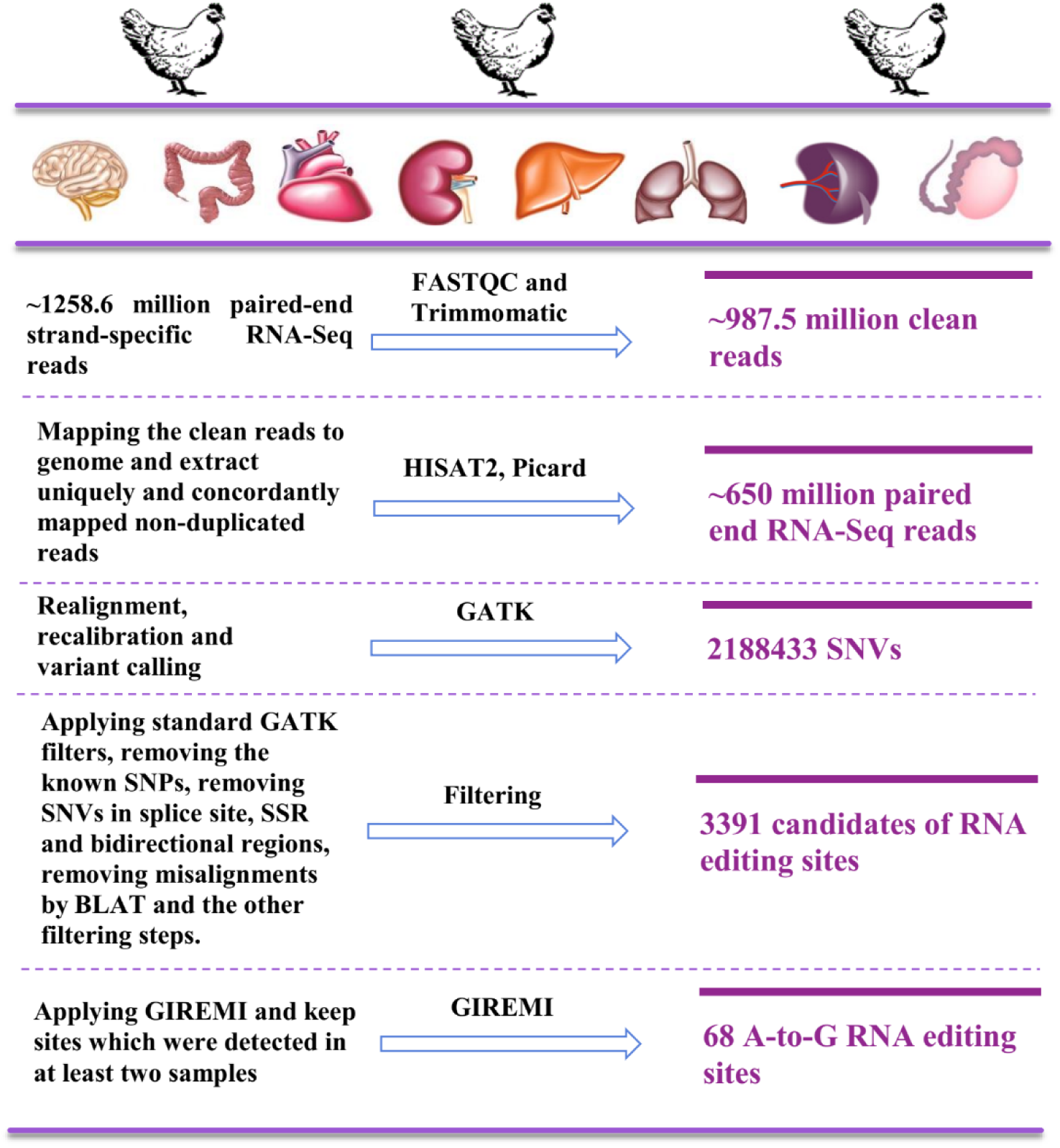
Simplified flowchart of our pipeline for RNA editing detection.

Totally, 1258.6 million pairs of raw reads were subjected to trimming and 987.5 clean reads were obtained. The clean reads were aligned to chicken genome with an average mapping rate of 78.48%. The rate of unique mapping into the reference genome was 81.16–94.30%. Summary statistics for reads and mapping information of different samples are provided in Supplementary File S1. After applying multiple filters with stringent thresholds, a total of 3391 candidate editing sites were identified across all samples (Figure 1). Finally, by using GIREMI algorithm, the number of candidate editing sites decreased to 84 sites. Also, these sites were detected in at least two biological replicates and covered by 10 or more independent reads, with at least three of them covering edited reads. As expected, the results revealed that A-to-G events were the most dominant type of modifications, which accounted for 81% (n=68) of all (Figure 2). Moreover, we found no C-to-U editing site. Therefore, we focused on the A-to-G editing sites for further analysis.

**Figure 2.**
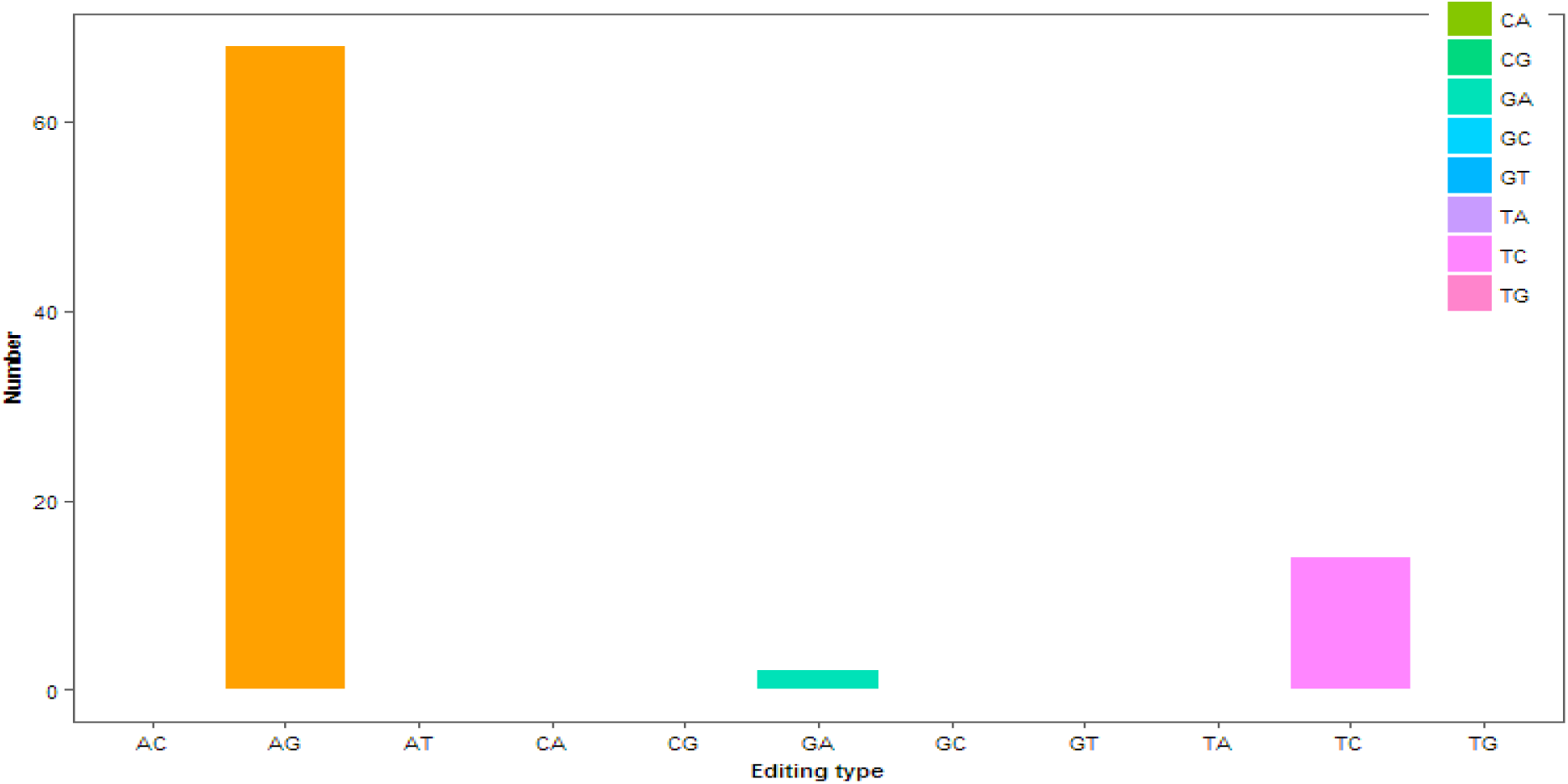
The number of each type of 12 possible nucleotide exchanges identified in all of the tissues.

### RNA editing distribution across tissues

We next investigated the distribution of A-to-G editing sites across tissues. We observed an uneven distribution of RNA editing sites across tissues. The highest number of A-to-G editing sites were identified in brain (41 sites), lung (37 sites) and heart (36 sites), respectively, which reflect the essential role of RNA editing in these tissues. Of the tissues profiled, the number of A-to-G editing site was lowest in liver tissue (8 sites). Of the 68 editing sites, 36 showed consistent altered editing in at least three tissues. Distribution of editing sites across tissues is displayed in Figure 3. Among all 68 the A-to-G editing sites, 42% had editing ratio <30%, whereas 17% had editing ratio >60%. In agreement with our previous study (Bakhtiarizadeh *et al*., 2018) in bovine, no significant difference was observed (using unpaired t-test and adjusted P-value <0.05) in the average editing ratio between different tissues. The average level of editing ratio was 0.38, 0.38 0.6, 0.40, 0.53, 0.37, 0.36 and 0.36 in brain, colon, heart, kidney, liver, lung, spleen and testes, respectively (Figure 4, Supplementary File S3). This range of editing ratio was in agreement with previous studies (Bazak *et al*., 2014, Picardi *et al*., 2015, Huntley *et al*., 2016). Furthermore, we found 36, 24, 21, 7, 28, 20 and 13 edited genes in brain, colon, heart, kidney, liver, lung, spleen and testes, respectively (Supplementary File S2).

**Figure 3.**
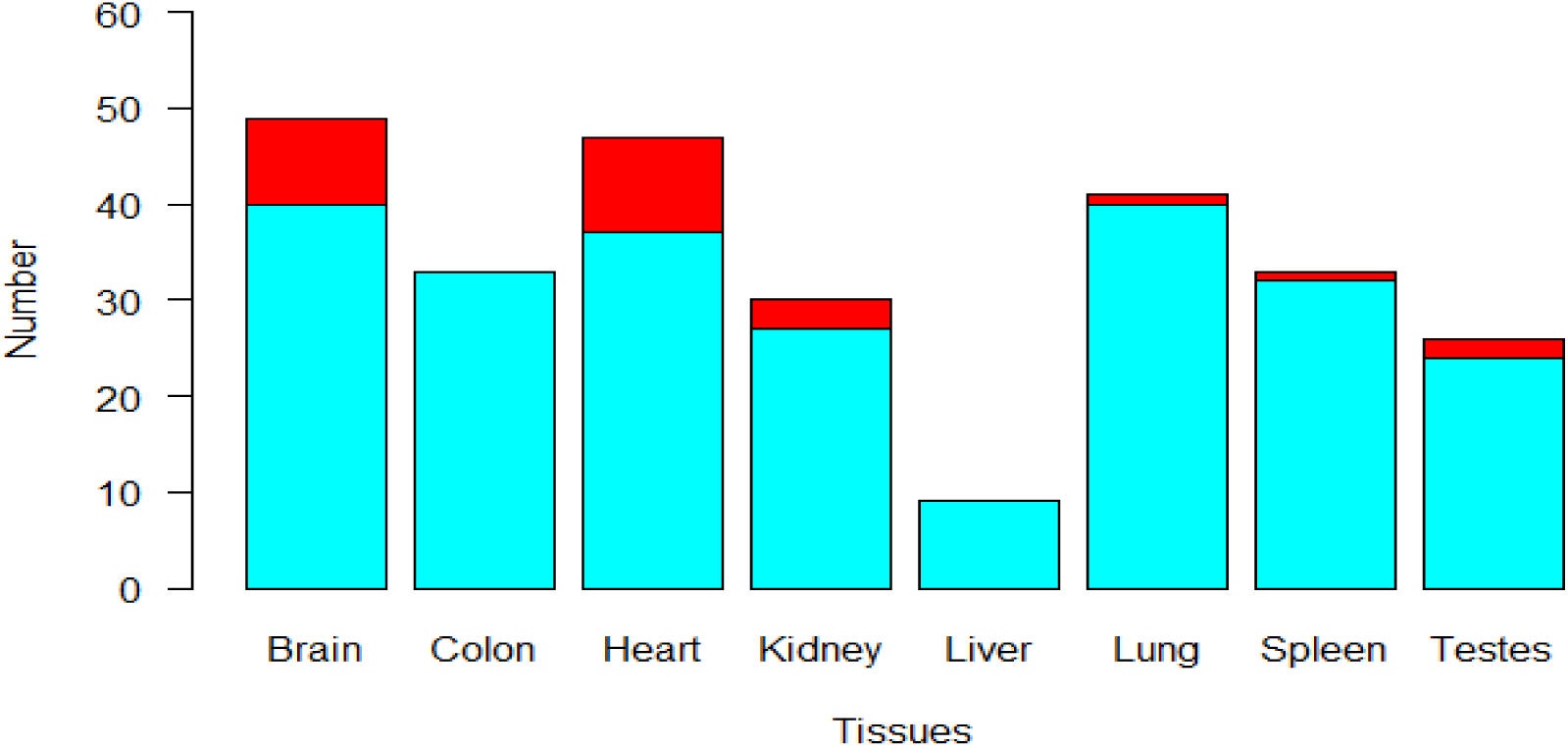
RNA editing distribution across chicken tissues. The red color represents the tissue-specific RNA editing sites.

**Figure 4.**
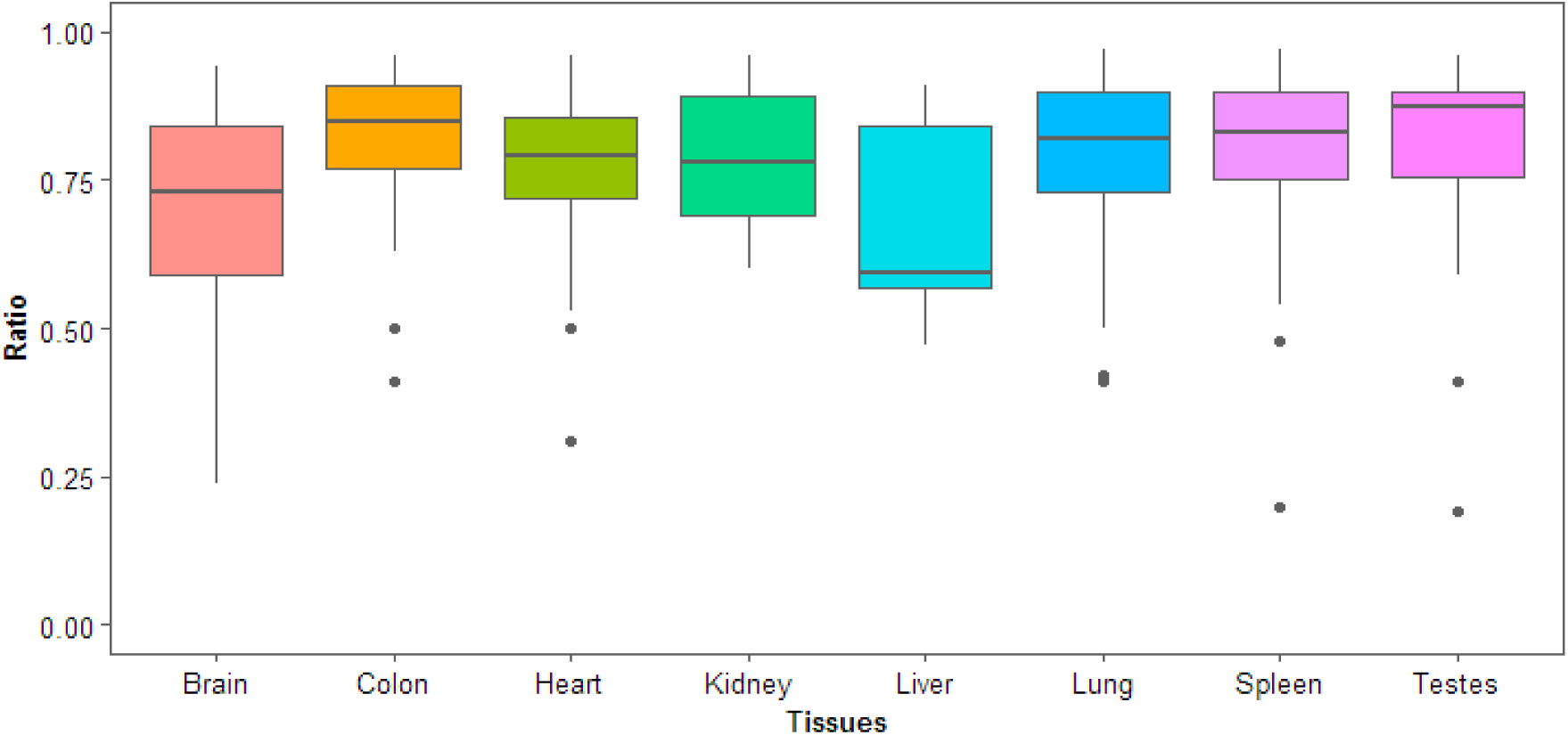
Editing ratio across different tissues.

### Distribution of RNA editing across different genomic regions

Among the 68 A-to-G editing sites, 34 and 23 sites were located within the known genes and their 5 kb flanking regions (16 sites in downstream and seven sites in upstream), respectively. Also, 11 of these editing sites occurred in the intergenic regions. Out of 34 editing sites in know genes, half of them resided in non-coding regions (eight sites in intron, eight sites in 3’ UTR and one site in 5’ UTR). Of the remaining 17 sites that were located in exons, 11 sites (32%) were identified as non-synonymous changes (with moderate effect), that resulted in amino acid recoding on nine genes (Table 1, Figure 5). Further concentration on the non-synonymous changes revealed that nine of 11 recoding sites were located at the first codon positions in protein-coding genes. Overall, the non-synonymous to synonymous ratio was 1.8, as expected, due to ADAR enzymes almost randomly edits mRNA targets (Maas *et al*., 2003). We observed that SLC30A5, ENSGALG00000038663, ENSGALG00000042073, ENSGALG00000046550 and ENSGALG00000044731 were non-synonymously edited in more than 50% of the tissues. Distribution of synonymous and non-synonymous editing sites in different tissues is shown in Figure 6.

**Table 1.** List of edited genes with non-synonymous effect

| Position | Gene | Effect of editing | Position | Gene | Effect of editing |
| --- | --- | --- | --- | --- | --- |
| 2: 39976218 | TGFBR2 | Isoleucin to Valin | 4: 60599161 | ENSGALG00000046550 | Threonine to Alanine |
| 3: 107399503 | XKR6 | Arginine to Glycine | 7: 4566825 | ENSGALG00000042073 | Arginine to Glycine |
| 4: 10844152 | GABRA3 | Isoleucin to Methionine | 12: 11517007 | ENSGALG00000038663 | Glutimine to Arginine |
| 19: 4827618 | GBAS | Serine to Glycine | 31: 20392 | ENSGALG00000044731 | Threonine to Alanine |
| Z: 21898144 | SLC30A5 | Isoleucin to Valine | 31: 20464 | ENSGALG00000044731 | Lysine to Glutamic acid |
| - | - | - | 31: 21035 | ENSGALG00000044731 | Asparagine to Aspartic acid |

**Figure 5.**
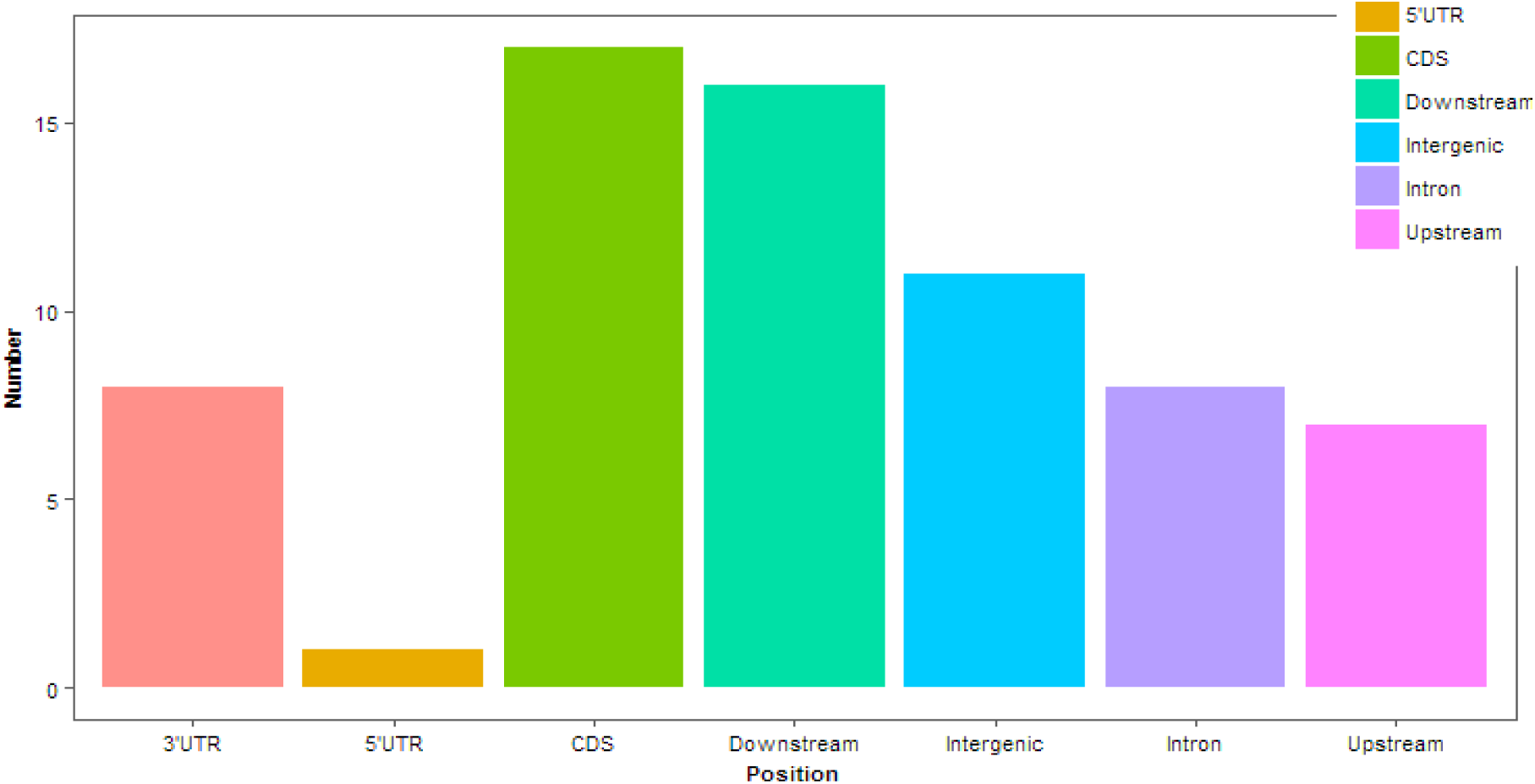
Distribution of RNA editing sites in genomic regions.

**Figure 6.**
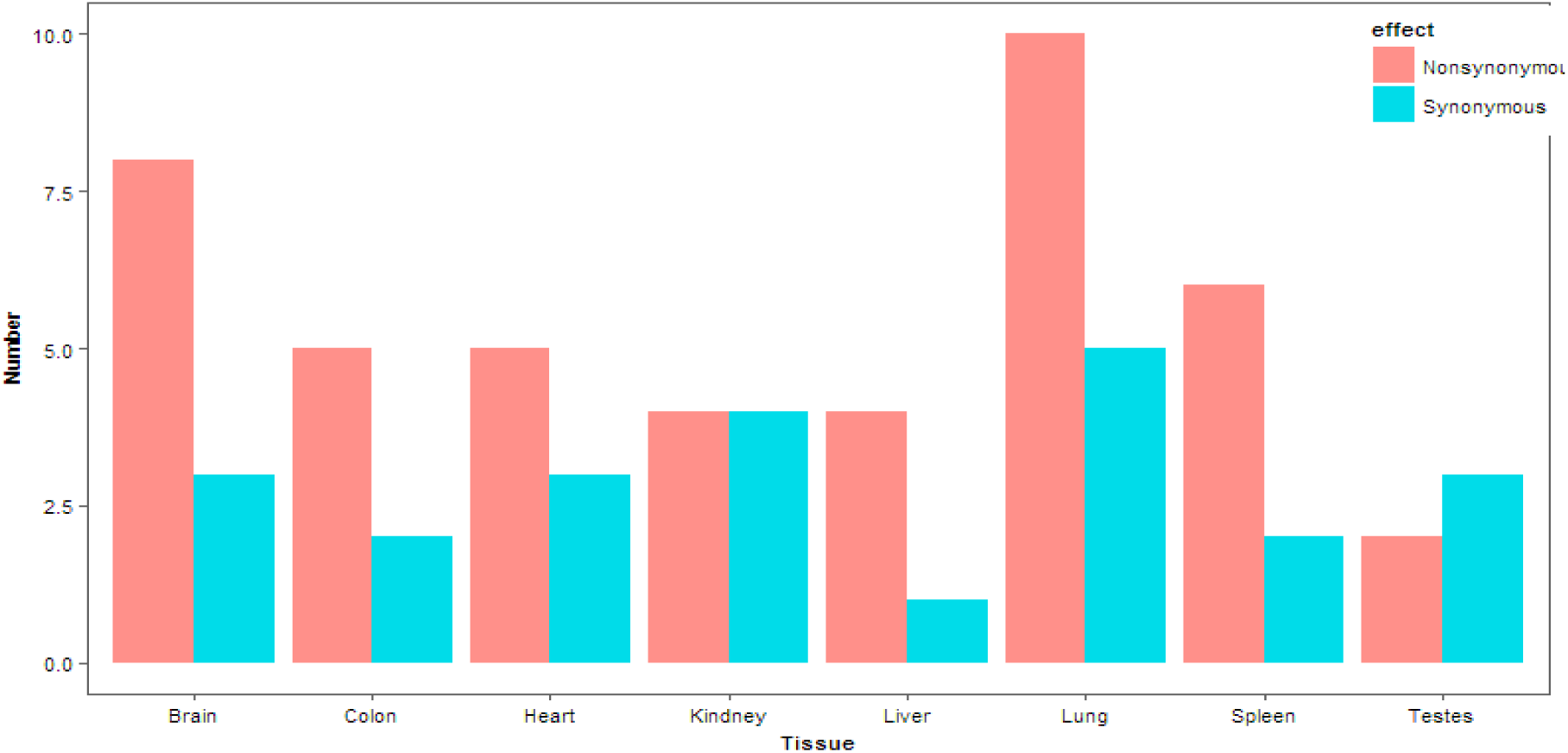
Numbers of synonymous and non-synonymous RNA editing sites in different tissues.

The average editing ratio of editing sites were estimated as 37% in intergenic, 46% in 5’ UTR (only one site), 35% in intron, 48% in non-synonymous, 42% in synonymous and 34% in 3’ UTR regions. Moreover, the average editing ratio of non-synonymous A-to-G editing sites were higher than that of synonymous editing sites in all of tissues (except kidney) (Supplementary File S3).

Since it is well known that A-to-G editing events are enriched in inverted repeat sequences (Daniel *et al*., 2014) we aimed to examine the enrichment of editing sites in these regions, in chicken genome. In contrast to mammals, we found that a few numbers of edited sites (five sites) resided within repetitive regions, such as LINEs and SINEs. Our genome wide screening showed that one, one and three edited sites were located in GGLTR7A, CHARLIE12 and CR1 family elements, which are related to SINE, DNA transposon and non-LTR retrotransposon classes, respectively (Supplementary File S4). A circos plot of the genomic landscape of the identified RNA editing sites is presented in Figure 7. Furthermore, positions of the edited genes and repetitive elements along with editing ratio of the sites are represented.

**Figure 7.**
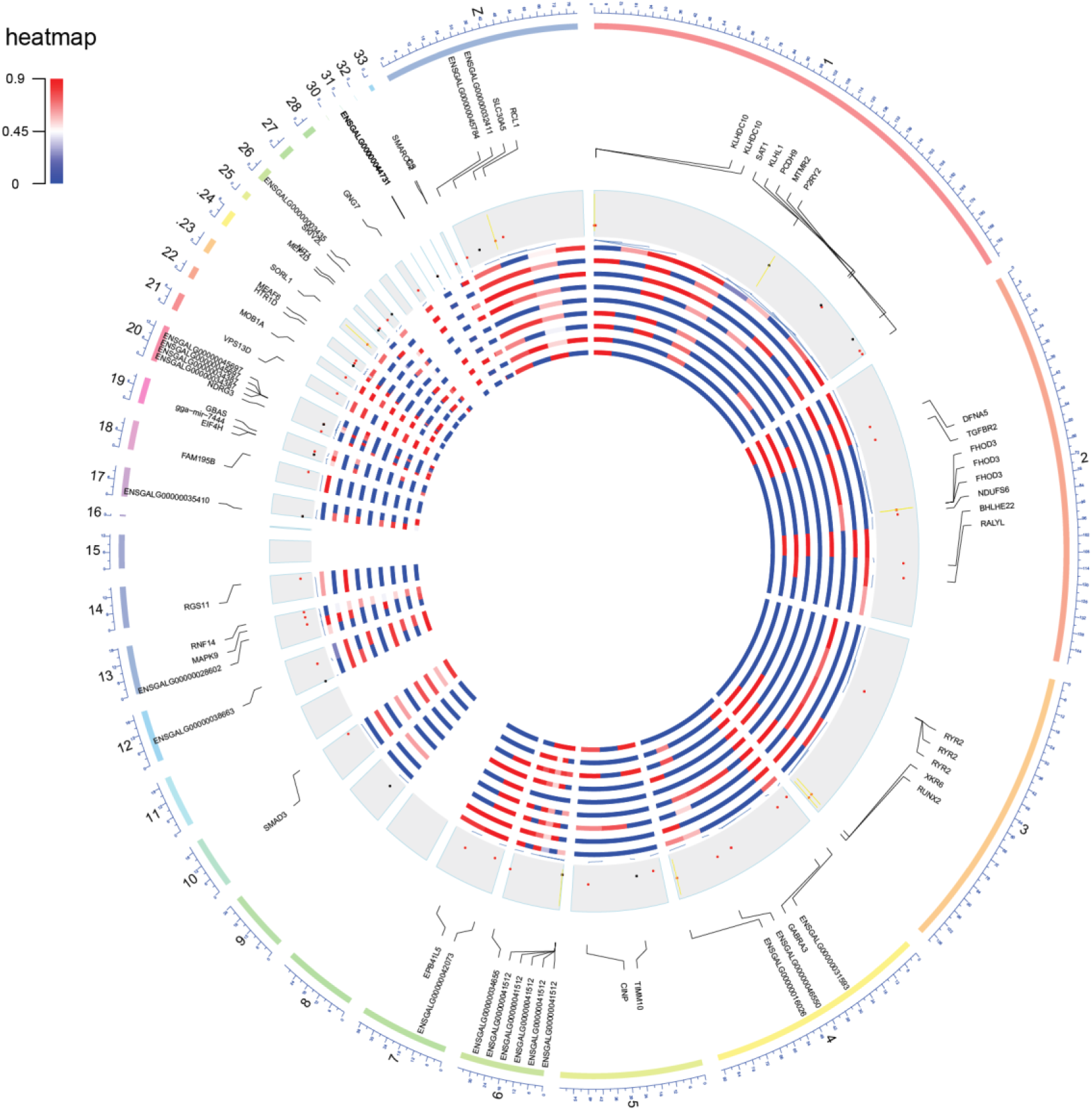
A Circos plot of the genomic landscape of the RNA editing sites in the chicken. The outermost ring shows chromosome numbers. The position of A-to-G editing sites (red circles) and non-A-to-G mismatches (black circles) are shown in the inner light gray rings. Also, the vertical yellow lines in the inner light gray rings represents the position of the repetitive elements (such as LINE, SINE and DNA transposons). The heatmap display the editing ratio of each sites in different tissues (tissues from the outside: brain, colon, heart, kidney, liver, lung, spleen and testes).

### Tissue-specific RNA editing

In addition to the differences in the editing ratio at different tissues, A-to-G editing sites appeared to be tissue specific. Hence, we concentrated on tissue specificity of the identified A-to-G editing sites and found a total of 20 A-to-G events which were edited exclusively or preferentially in only one tissue. The highest number of tissue specific editing sites was detected in brain (eight sites). Also, heart (seven sites), kidney (three sites), lung (one site) and testes (one site) were the other tissues with tissue specific editing sites. These tissue specific sites were located on six, four and one genes in brain, heart and lung, respectively. Moreover, 12 sites were found on nine genes to be shared between two tissues. Also, we found 11 sites on nine genes, including RNF14, SLC30A5, ENSGALG00000045784 and ENSGALG00000041512, which were edited in at least six tissues. The A-to-G editing sites that were unique or common among different tissues are provided in Supplementary File S3.

### ADAR expressions

There are two members of ADAR family in the chicken, including ADARB1 (ADAR2) and ADARB2 (ADAR3). Expression of these genes varied across the tissues, as ADARB1 was uniformly expressed in all tissues (the highest expression observed in brain). As with mammals (Picardi *et al*., 2015, Picardi *et al*., 2017, Bakhtiarizadeh *et al*., 2018), ADARB2 was exclusively expressed in brain (Figure 8). In spite of the questionable editing activity of ADARB2 (CHEN *et al*., 2000), this result has raised the suspicion that the expression pattern of this gene can explain why more A-to-G editing sites occurred in the brain than the other tissues. Therefore, ADARB2 may have the tissue specific activity to edit genes in the brain.

**Figure 8.**
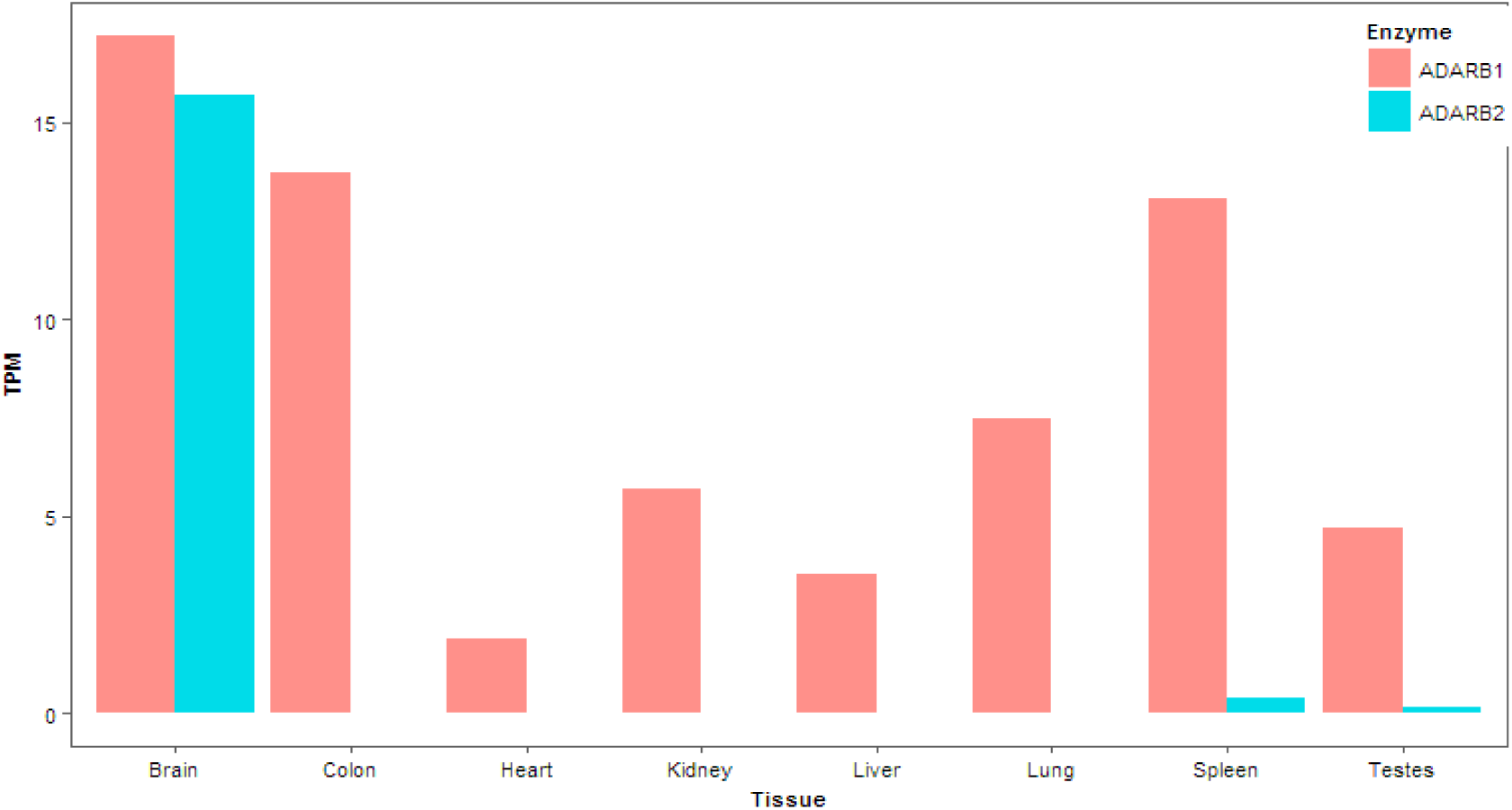
Expression levels of ADAR enzymes across different tissues.

We then investigated Spearman’s correlation between a) ADARB1 expression and editing ratio, b) ADARB1 expression and editing number, c) ADARB2 expression and editing ratio and d) ADARB2 expression and editing number, in all of the tissues. We applied this analysis to determine whether the variation in editing ratio/numbers in different tissues can be explained by the expression of ADAR enzymes. The results showed a positive correlation, but not significant, between ADAR enzymes expression and editing ratio/numbers in all of the comparisons (cor=0.31 and p<0.46 for ADARB1 and editing ratio, cor=0.37 and p<0.37 for ADARB1 and editing numbers, cor=0.12 and p<0.77 for ADARB2 and editing ratio, cor=0.25 and p<0.55 for ADARB1 and editing numbers). We also checked Spearman’s correlation between the expression levels of the edited genes and their editing ratio in all tissues to investigate whether expression levels of edited genes are associated with RNA editing levels in each tissue. The correlation coefficients ranged from −0.76 to 0.87 (Supplementary File S5).

### EST analysis

To validate our results, we used the public EST chicken sequences and investigated whether there are any EST sequences containing the identified editing sites. Out of 68 editing sites, 26 sites (38%) were found in at least one edited EST sequence (Supplementary File S4).

### Sequence preferences analysis

The nucleotides neighboring the identified A-to-G editing sites showed a clear pattern consistent with the known ADAR sequence preference (Figure 9). According to previous studies (Danecek *et al*., 2012, Bazak *et al*., 2014, Picardi *et al*., 2015), we observed an under- and over-representation of guanosine immediately upstream and downstream (relative location 1) from the edited nucleoside.

**Figure 9.**
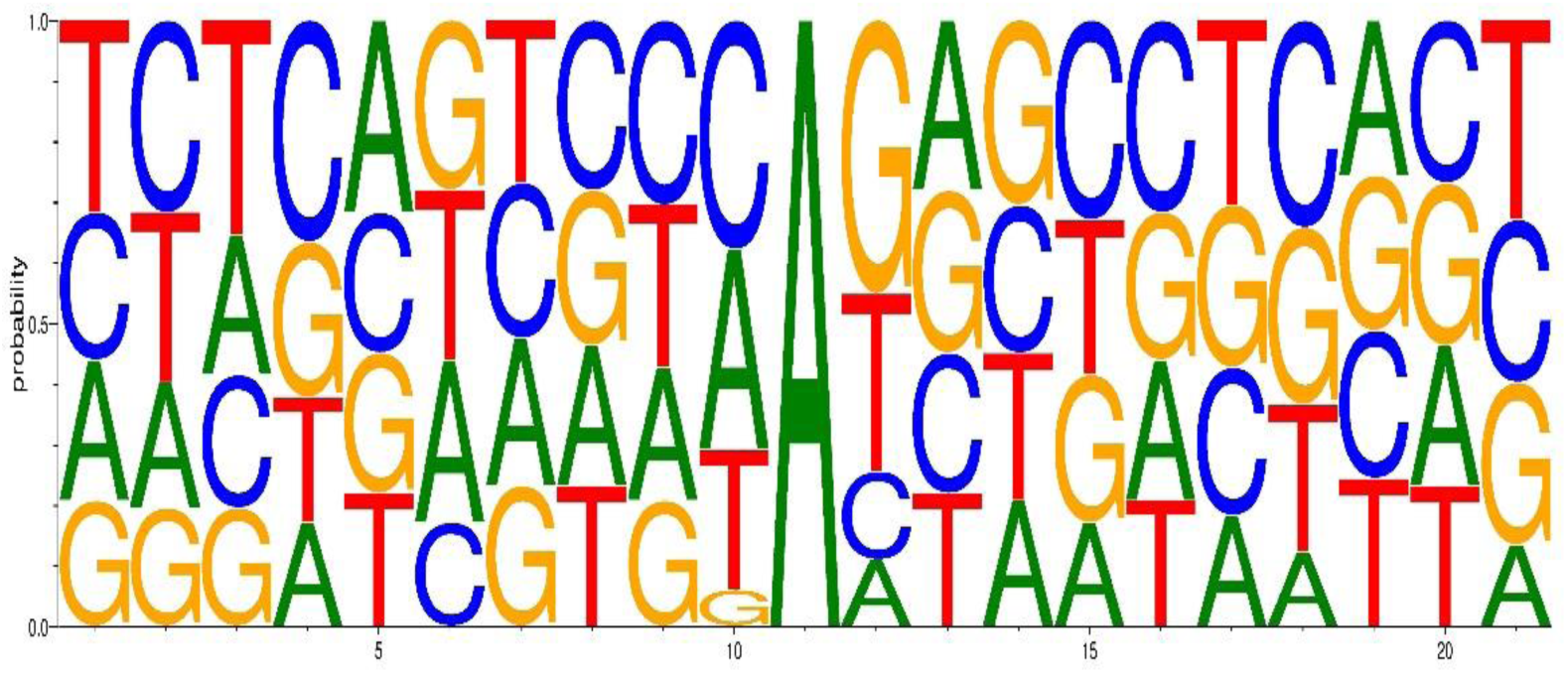
Sequence preferences of the neighborhoods of A-to-G RNA editing sites.

## Discussion

To investigate how A-to-G editing sites are distributed across different chicken tissues and if there is any C-to-U editing event, we analyzed a large number of strand-specific RNA-Seq data from public database. The first challenging issue to address for discovering RNA editing sites is to correctly align short reads to the reference genome and mitigate as much as possible the effect of spurious read alignments. This issue may lead to not only inaccurate estimates of RNA editing sites but also false-positive predictions of editing events (Kim *et al*., 2015). Here, we used Hisat2, as the efficiency of this aligner is already highlighted in aligning the reads to genome for predicting RNA editing sites(Xiong *et al*., 2017, Yao *et al*., 2017). In addition, we used a stringent filtering method after mapping, including extraction of uniquely and concordantly mapped reads, removing of the duplicated reads, realignment around putative insertions and deletions and recalibration of base quality values. Furthermore, the effect of misalignments was minimized by using BLAT, a more accurate aligner than fast-mappers like Hisat2, on reads with mismatches. It is highly desirable to have a method that identify editing sites without the need to filter out SNPs. Following aligning the reads, we identified candidate RNA editing sites, by means of a stringent computational procedure requiring only RNA-Seq data and no a priori knowledge of genomic reads from the same sample. In this regard, GIREMI enabled us to filter false positives sites due to polymorphic variations in the genome. Furthermore, to distinguish more between SNPs and RNA editing events, we made the assumption that editing sites must be present in at least two biological replicates. Hence, a strong RNA editing site is required to be appeared in least two biological replicates, whereas false positives results (such as SNPs) are unlikely to be common to unrelated individuals.

Genome-wide identification of RNA editing in all eight tissues yielded a total of 84 potential editing events. In this study, the results revealed a clear enrichment for A-to-G changes over all other changes (~81%), indicating that the results are reliable. Similar to what has been reported in chicken (Frésard et al., 2015, Roux et al., 2016), we also observed no C-to-U editing site, indicating that this type of editing mechanism is missing in the chicken. An obvious explanation for the lack of observed C-to-U editing events in chicken would be lack of a homolog of APOBEC1 enzyme in the chicken genome (Conticello *et al*., 2004). On the other hand, C-to-U editing is a very rare event in mammals (Blanc and Davidson, 2010, Bakhtiarizadeh *et al*., 2018). This point reinforces the hypothesis that A-to-G is the only RNA editing event in chicken and is restricted to ADAR-mediated events. However, further studies on the current topic are required in order to confirm the hypothesis. The second and third type of the identified events were T-to-C and G-to-A mismatches. Interestingly, similar results are reported in human (Ramaswami *et al*., 2013, Franzén *et al*., 2018) and bovine (Bakhtiarizadeh *et al*., 2018), which used strand-specific RNA-Seq data for RNA editing detection. On the other hand, we can use the ratio of G-to-A to A-to-G mismatches as a criterion for evaluating the specificity of the RNA editing detection, as have been used in previous studies (Levanon *et al*., 2004, Bahn *et al*., 2012, Liscovitch-Brauer *et al*., 2017). This assumes that G-to-A events in strand-specific RNA-Seq data are sequencing or mapping errors (Levanon *et al*., 2004). Hence, the false discovery rate of our study is low at 3% (2/68), further corroborating the authenticity of our results.

To our knowledge, it is the first in-depth study to investigate the RNA editing patterns across more than two chicken tissues. Our results showed that the extent of A-to-G editing is limited (68 sites) in different chicken tissues, a finding in good agreement with previous studies on chicken (Frésard et al., 2015, Roux et al., 2016). One possible explanation for this result is lack of ADAR1 enzyme in chicken. Another explanation for the low number of editing sites in chicken is low frequency of interspersed repeats in the genome of this species. It is well known that ADAR enzymes bind to double-stranded structures and perform A-to-G editing. Hence, any transcript that forms long double-stranded structures, such as LINE and SINE families, can be suitable as an ADAR target (Reenan, 2001). We found A-to-G editing sites were weakly enriched in interspersed repeats (SINE, LINE and DNA transposons), which is consistent with low number of these repeats in the chicken genome. For example, while interspersed repeats comprise around half of the mammalian genome, only 10% of the chicken genome is comprised of these repeats (Lee et al., 2017).

Our findings showed that the identified A-to-G editing sites shared the similar characteristics with those of previous studies in mammals as well as in chicken. The characteristics can be summarized as follow:

1. Both nucleotides surrounding A-to-G editing sites can affect the specificity and efficiency of editing, as it might possibly facilitate the binding of ADARs to their targets (Eggington *et al*., 2011). Although the mechanisms of target recognition by ADAR enzymes are not well understood, it has been shown that under- and over-represented nucleotide in the 5’ −1 and the 3′ +1 of A-to-G editing sites is guanine in mammals (Danecek *et al*., 2012, Bazak *et al*., 2014, Picardi *et al*., 2015). Here, when the flanking sequences of all A-to-G editing sites were analyzed, a similar sequence-preference between chicken and mammals was observed (Bazak *et al*., 2014, Picardi *et al*., 2015, Huntley *et al*., 2016, Bakhtiarizadeh *et al*., 2018), suggesting similar ADAR preference among mammals and birds (Bazak *et al*., 2014, Picardi *et al*., 2015). Interestingly, under-representation of A nucleotide in regions upstream and downstream editing sites is consistent with other studies in human (Picardi *et al*., 2015) and rhesus macaque (Chen *et al*., 2014).
2. In our study, 38% of the A-to-G editing sites were confirmed by EST sequences. The number of validated editing sites by EST sequences was lower than the previous studies in bovine (Bakhtiarizadeh *et al*., 2018) (with 66% of validation) and mouse (Lagarrigue *et al*., 2013) (with 58% of validation). It can be attributed to significantly lower number of EST sequences of chicken in NCBI database, as there are only 600,435 ESTs in chicken in comparison with 1,583,417 and 4,871,060 ESTs in bovine and mouse, respectively.
3. Similar to previous studies in chicken (Frésard et al., 2015, Roux et al., 2016) and in contrast to mammals (Yablonovitch *et al*., 2017), we found that both the frequency and editing ratio of non-synonymous A-to-G editing sites were higher than those of synonymous editing sites (Supplementary File S3). This finding suggests that RNA editing mechanism may be important for proteome diversity in chicken. It may have important effects on protein translation, which has also been reported in human (Picardi *et al*., 2015).
4. Here, in consistent with previous studies in mammals (Picardi *et al*., 2015, Huntley *et al*., 2016, Roux *et al*., 2016, Bakhtiarizadeh *et al*., 2018) as well as in chicken (Frésard *et al*., 2015, Roux *et al*., 2016), A-to-G editing sites mainly occurred in non-coding regions (Figure 6). Also, abundance of the identified editing sites on the non-coding sequences were similar between our study and the other studies on chicken, as the editing sites were enriched on downstream, intergenic and intronic regions, respectively (Frésard *et al*., 2015, Roux *et al*., 2016). Since, annotation of the flanking regions of genes are hard and can also vary in size (Jan *et al*., 2011), some of the downstream or upstream regions can be considered as extended 3’ or 5’ UTR of genes, respectively. Further analysis of the editing sites showed that these sites were more frequently in 3’UTRs (12%) than in 5’UTRs (1%), which is similar to observations in humans (Peng *et al*., 2012a), bovine (Bakhtiarizadeh *et al*., 2018), drosophila (St Laurent *et al*., 2013) as well as in chicken (Frésard *et al*., 2015) and highlight the importance of A-to-G editing in 3’ UTR region. Therefore, in line with previous studies (Venø *et al*., 2012, Picardi *et al*., 2015, Huntley *et al*., 2016, Bakhtiarizadeh *et al*., 2018), our results suggest a potentially close relationship between editing events in non-coding regions and regulation of gene expression. For example, we found eight editing sites in intron, which may contribute to mRNA alternative splicing regulation (Higuchi *et al*., 1993). Also, eight of editing sites were located in 3’UTR (a target region for transcript regulation), which can alter mRNA stability and affect miRNA binding (Zhang *et al*., 2017). Hence, A-to-G editing events might play an important role cooperating with miRNAs in regulating gene expression. However, the function of non-coding RNA editing is an open area of research and further investigations are needed to understand the functional effect of these events.
5. To identify the potential molecular mechanisms underlying the different RNA editing activities across tissues, we investigated the different hypothesizes. First of all, we analyzed the Spearman’s correlation between ADAR enzymes expressions and the levels of editing ration/numbers in different tissues. Although, none of the correlations were significant, there were some trend in the correlations between editing ratio/numbers and ADAR expression. In other words, a small part of the present differential editing across tissues can be explained with ADAR enzymes expression, that were not expressed uniformly in all tissues. An alternative hypothesis explaining differential RNA editing levels across tissues is that expression levels of edited genes are associated with RNA editing levels meaning that the increased expression of a given gene can lead to higher editing levels. In testing this hypothesis, we observed no meaningful pattern between the expression levels of the edited genes and their editing ratio across tissues. These findings confirmed previous works that have shown this trend in human (Picardi *et al*., 2017, Yao *et al*., 2017) and cattle (Bakhtiarizadeh *et al*., 2018). Based on some of the previous studies (Huntley *et al*., 2016, Bakhtiarizadeh *et al*., 2018), it is more possible to derive a clear relationship between editing ratio of tissues specific edited genes and their expressions. Hence, to assess this hypothesis, we focused on tissue specific editing sites on the edited genes. The results revealed that seven out of 11 tissue specific edited genes had the highest expression in the tissue that they were edited (Supplementary File S6). This finding at first glance implies a relative relationship between the editing ratio of tissues specific edited genes and their expressions. However, generally and in line with previous studies in mammals (Picardi *et al*., 2015, Huntley *et al*., 2016, Bakhtiarizadeh *et al*., 2018), we found no obvious pattern between ADAR enzymes or edited genes expression and RNA editing activities across chicken tissues. Therefore, more work remains to be done to refine our understanding of potential molecular mechanisms underlying the RNA editing activities across chicken tissues.

Generally, most interest has been focused on non-synonymous (recoding) editing events, where RNA editing results in amino acid changes. In this study, nine edited genes were found with non-synonymous changes. Their editing ratio (on average) ranged from high (83% of ENSGALG00000044731) to low (17% of ENSGALG00000042073), indicating that both edited and unedited genes are required for the proper function of the tissues (Supplementary File S3). Interestingly, we found that eight of the recoding editing events occurred in the brain, such as gamma-aminobutyric acid A receptor α3 subunit (GABRA3), glioblastoma amplified sequence (GBAS) and transforming growth factor, beta receptor II (TGFBR2) (Supplementary File S3). Of these, GABRA3 was exclusively edited in the brain with isoleucine to methionine change, which can lead to a change in the third transmembrane region (Daniel *et al*., 2011). This gene is a part of the major inhibitory neurotransmitter system (Nicholas *et al*., 2010) and express during development of the brain (Xiang *et al*., 2000). Also, it has been shown that GABRA3 is targeted by both ADARs enzymes (Ohlson *et al*., 2007). Interestingly, the editing site on GABRA3 was already reported in different animals such as human (Nicholas *et al*., 2010, O’Neil *et al*., 2017), mouse (Daniel *et al*., 2011), pig (Venø *et al*., 2012), bovine (Bakhtiarizadeh *et al*., 2018), rat, rabbit, dog, frog (Daniel *et al*., 2011) and chicken embryos (Daniel *et al*., 2011, Frésard *et al*., 2015). This result suggests that editing events may contribute to amino acid functional conservation throughout the evolution of vertebrates, as has been reported in previous study in chicken (Roux *et al*., 2016). Another gene that had recoding site was GBAS, which was edited in brain, heart and lung tissues. The enriched expression of this gene is reported in brain and heart (Smits *et al*., 2010). It has been revealed that GBAS plays important role in malignant form of central nervous system tumor. Also, mutation in this gene can lead to combined deficiencies in oxidative phosphorylation (Chen *et al*., 2011). Moreover, GBAS is introduced as a candidate gene in mitochondrial disorders. On the other hand, heart and brain have high energy demands and have been found to be often affected in mitochondrial diseases (Smits *et al*., 2010). We observed that the A-to-G editing events on TGFBR2 occurred in four tissues including brain, colon, lung and spleen, with the highest editing ratio in the brain. This gene encodes a cell-surface transmembrane receptor for transforming growth factor-ß (TGF-ß). The predominant expression of TGFBR2 has been shown by neurons in the central nervous system (Lim *et al*., 2011). Also, it has been reported that mutations in this gene are very common in glioblastoma, most aggressive malignant primary brain tumor in humans (Sivadas and Kannan, 2014). Therefore, functions of the edited genes in the brain corroborated the relevant biological role of these genes with nervous system. This is compatible with most previous studies, which reported that RNA editing is enriched in the central nervous system and plays specific roles in brain physiology (Li and Church, 2013, Picardi et al., 2015, Huntley et al., 2016).

Tissue specificity of some RNA editing sites have been postulated in several studies and in different species (Alon *et al*., 2015, Picardi *et al*., 2015, Funkhouser *et al*., 2017, Bakhtiarizadeh *et al*., 2018), especially in chicken (Frésard *et al*., 2015). Here, we specifically looked for tissue specific A-to-G editing sites. In consistent with previous studies (Picardi *et al*., 2015, Picardi *et al*., 2017, Bakhtiarizadeh *et al*., 2018), brain showed the highest number of tissue specific editing sites along with the highest number of tissue specific edited genes. An interesting aspect emerging from our results was the finding of three (GABRA3, sortilin-related receptor 1 (SORL1) and serotonin receptors 1D (HTR1D)) and two (cardiac ryanodine receptor (RYR2) and formin homology 2 domain containing 3 (FHOD3)) tissue specific edited genes in brain and heart, respectively, with completely related functions. It is well documented that SORL1 is genetically associated with Alzheimer disease (Rogaeva *et al*., 2007). In addition, down-regulation of this gene in the brain of patients has been suggested a causal role for SORL1 in the pathogenesis of Alzheimer. We found a synonymous editing site in coding sequence region of HTR1D, which encodes a serotonin receptor, and is thought to be involved in the development of migraine (Ruaño *et al*., 2007). This gene is reported as responsible for regulating the release of serotonin in brain (Xiang *et al*., 2008). Also, previous studies have found associations between polymorphisms in this gene and anorexia nervosa (Woutersen, 2013). These results further highlighted the importance of RNA editing in the brain. As aforementioned, RYR2 and FHOD3 were found as tissue specific edited genes in heart with three and two editing sites in their introns. RYR2 encodes a calcium ion transporter in the sarcomeric reticulum and its function in the heart is well known (Wehrens *et al*., 2004). This gene is required for excitation-contraction coupling in the heart to keep the receptor closed during the resting phase of the heart. It is reported that dysfunction of RYR2 is associated with heart failure (Marks, 2001). FHOD3 is a cardiac member of the formin family proteins, which has been demonstrated to function in heart development (Rosado *et al*., 2014). This gene is highly expressed in the heart and is necessary for myofibrillogenesis during cardiac development, especially in the maturation of myofibrils (Kan-o *et al*., 2012). These results indicate that A-to-G editing may not only be of importance for nervous system, but also be involved in regulating the functions of genes in heart and probably in other tissues. Therefore, edited genes (especially tissue specific ones) in all of tissues deserve careful attention and further studies.

To further evaluate our results, we compared the A-to-G RNA editing sites with those identified in the three previous studies in chicken (Frésard *et al*., 2015, Roux *et al*., 2016, Hung *et al*., 2017). Since the coordinates of our editing sites were based on Galgal5, liftOver program (http://hgdownload.cse.ucsc.edu/admin/exe/) from UCSC was applied to convert genomic coordinates from Galgal4 (in other studies on chicken) into Galgal5. We only identified one site on GABRA3 in common with (Hung *et al*., 2017) and one site on upstream region of ENSGALG00000034655 gene in common with (Frésard *et al*., 2015). We also observed low overlap among the three studies in chicken (Supplementary File S7). The limited overlap can be attributed in large part to differences in computational strategy, experimental design, read length and variability in the sequencing type (single/paired-end or stranded/non-stranded) as well as sequencing depth of samples, whereas the importance of these factors is highlighted in previous studies (Diroma *et al*., 2017, Franzén *et al*., 2018). For example, Hung et. al. had been used a low coverage (~11 million reads per sample in average), non-strand-specific and single-end RNA-Seq data on six different chicken tissues, including cerebellum, brain, kidney, liver, heart and testes. Also, the two other studies applied non-strand-specific RNA-Seq data. However, we used strand-specific and paired-end RNA-Seq data with sufficient coverage on eight tissues. In the studies that use non-strand-specific RNA-Seq data, all of T-to-C changes are considered as canonical editing events (A-to-G). On the other hand, it is well known that many of these sites are not real A-to-G editing sites (Ramaswami *et al*., 2013).

## Conclusions

Here, we used 22 samples from three chickens to provides a detailed picture of RNA editing across eight chicken tissues. RNA-Seq samples were analyzed using a novel and rigorous computational pipeline. From a bioinformatic point of view, our results highlighted the importance of properly filtering for bias in RNA-Seq datasets along with the necessity of using biological replicates. From our analysis, 68 A-to-G and no C-to-U editing sites were identified on chicken. The identified A-to-G events constituted 66 novels and two previously reported editing sites. Also, only 16% of the A-to-G editing sites were protein recoding events, indicating that protein recoding may not be the primary function of editing mechanism in chicken. This study represented a comprehensive list of RNA editing events in eight chicken tissues. Now, we can begin to determine the functional importance of editing, whether editing events might differ between the tissues.

